# A traveling network model predicts emergent dynamics and search behavior from local remodeling in *Physarum polycephalum*

**DOI:** 10.64898/2026.08.13.744445

**Authors:** Arnold Chen, Shenghao Tan, Yash V. Mundewadi, Ingmar H. Riedel-Kruse, Nate J. Cira

## Abstract

A variety of connected systems, ranging from the cytoskeleton to human organizations, dynamically rearrange themselves in order to move through physical or abstract space. However, our understanding of how systems-level behaviors arise from local restructuring actions remains limited, necessitating comparison of real-world data to models that predict network structure and dynamics. To understand these systems, we study an accessible example, the branching slime mold *Physarum polycephalum*, by imaging the organism as it travels and extracting key fundamental quantities from its continuously remodeling tubular network. By using these quantities as input parameters to a traveling network model, we find that with no further fitting, the model quantitatively matches key emergent properties from *P. polycephalum* dynamics including path length, relocation time, and search efficiency at different spatial resolutions. These findings demonstrate how a traveling network model can capture *P. polycephalum* behaviors, highlighting the potential to use traveling networks more broadly for understanding and predicting connected dynamic systems by linking local measurements to emergent, system-wide behaviors.

## Introduction

The slime mold *Physarum polycephalum* exhibits complex behaviors and morphologies despite lacking a brain or nervous system. These behaviors include memory-encoding (1–4), the ability to anticipate environmental changes or stimuli (5,6), and the ability to solve optimization-related problems such as finding a path through a maze and connecting food sources in a way that resembles features of an existing railway network (7,8). Specifically, insights from *P. polycephalum* optimization have been applied towards the Steiner Tree Problem to minimize edge weights in a tree (9), ant colony optimization algorithms (10), and traffic assignment and optimization (11,12).

Numerous models have been employed to understand *P. polycephalum* behavior. These have focused on different aspects of the organism such as tube fluxes (8,13,14), memristive behavior (5), and network growth (15). Other models focus on the overall network pattern, including occupying positions in fixed lattices (16,17). Some point source or network models exhibit less regular morphologies (18–20). Past network models typically focus on how the organism expands through space and change its structure to optimize specific connectivity outcomes. One key behavior that is seldom modeled is the organism’s ability to travel spatially away from its initial location. This is important to understanding how the organism searches and navigates its environment over time. Past attempts to understand how the organism travels through space have quantified the diffusion of the center of mass of the organism (20,21), but do not simultaneously capture the full dynamic network structure.

Recently, we introduced the concept of a “traveling network” along with a model that considers how dynamic networks travel through space as the result of restructuring (22). This model explicitly tracks the full network structure and dynamics, which makes it possible to see how restructuring actions result in different system morphologies and behaviors. A traveling network framework applies to other systems, well beyond slime molds. For instance, the actin cytoskeleton depolymerizes and elongates which affects the higher-level search behavior of the cells (23,24). Swarm robots restructure their communication tree to achieve highly complex collective behavior (25). Companies traverse through the space of market opportunities, such as seeding, nurturing, and pruning to increase overall productivity (26). Among these examples, *P. polycephalum* stands out as especially tractable for experimental study, enabling us to observe and measure both local restructuring and system-level behaviors and potentially allowing elucidation of general principles applicable to this class of systems.

While the theoretical underpinnings of a traveling network model were rigorously developed (22), grounded comparison to real-world data was limited, and it remains unclear how well any real system can be modeled as a traveling network. Here, we bridge this gap with the first comprehensive comparison of experimental data to results from a traveling network model. To do this, we recorded *P. polycephalum* behavior, extracting fundamental quantities, then we used these extracted quantities as network model input parameters, assessing how well the model captures key emergent behaviors of the organisms. We then modify the model based on observations to better describe specific search capabilities of the organism.

## Results

### Measurements of Physarum polycephalum’s varied branching and growth morphologies can be mapped to a traveling network model

*P. polycephalum* is a slime mold that displays branched morphology in the plasmodial stage of its life cycle (Fig. 1a). To record *P. polycephalum* behavior, we imaged the movement of the organisms on agar surfaces in petri dishes over multiple days. During this time, organisms move through the environment, moving away from their starting locations (Fig. 1b). From these recorded behaviors, we extracted various fundamental quantities including organism size, branching angle, retraction speed, and the number of branching events (Fig. 1c) (see methods).

**Figure 1:**
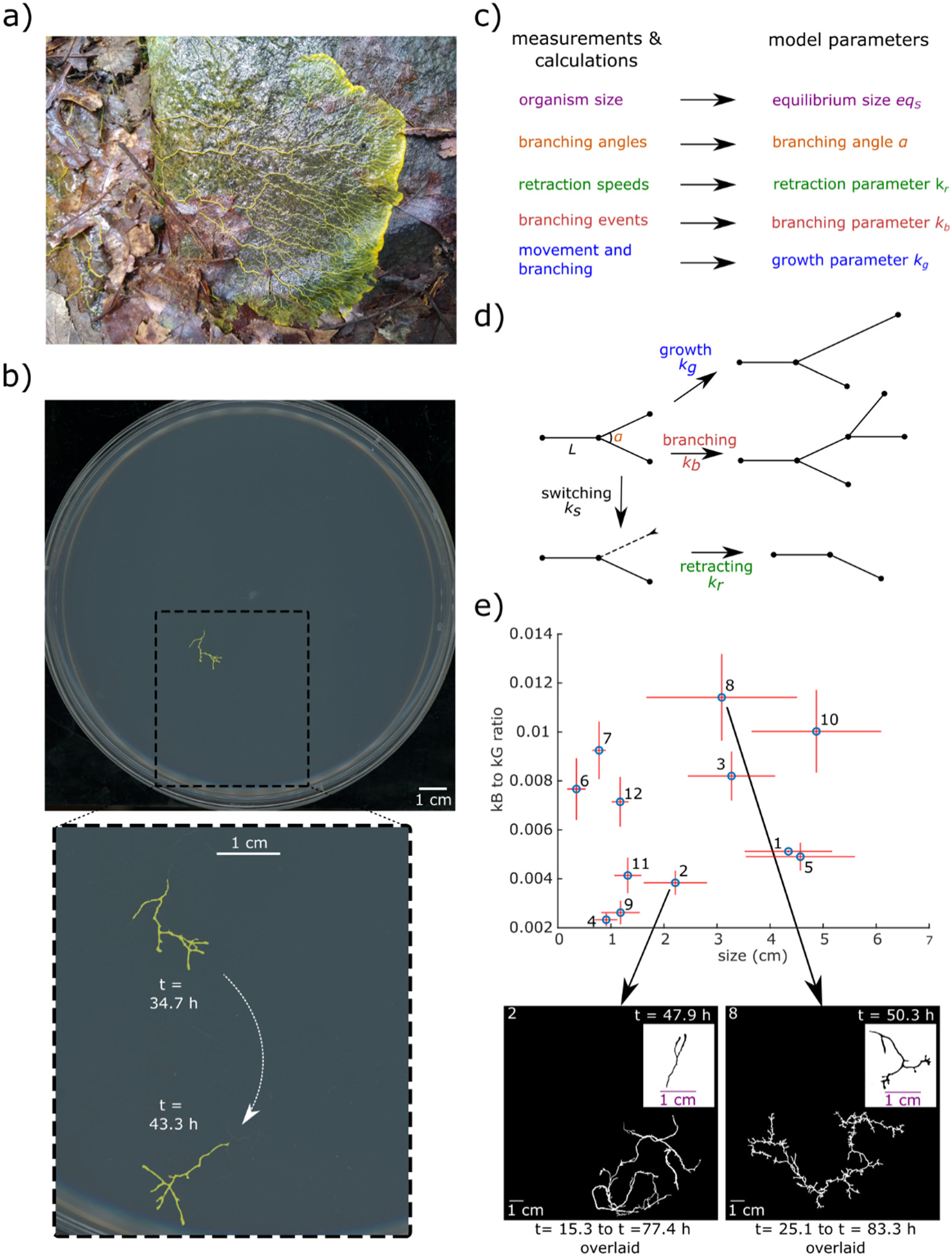
Time-lapse imaging of *P. polycephalum* can be mapped to a traveling network model. a) *P. polycephalum* is a slime mold with branched morphology. b) *P. polycephalum* travels spatially over time. Individual images show that the organism moved downwards from t = 34.7 h (top) to 43.3 h (bottom). Inset shows a zoomed-in overlaid image of the two individual images to illustrate that the entire organism has displaced in position through time. Color of the organism has been enhanced to better visualize against the background. c) Quantities were extracted from *P. polycephalum*, including average organism size, retraction speed, number of branching events, and branching angles. These are converted correspondingly to inputs for the model. d) A traveling network model and possible actions of the simulated organism (adapted from (22) fig. 1d). The branching angle is *α*, and the edge length is Δ*L*. The network branches at rate *k_b_*, grows at rate *k_g_*, and switches at rate *k_s_*. After switching, the edge retracts at rate *k_r_* until the edge is eliminated. e) The amount of branching relative to growth, represented by the ratio between the parameters *k_b_* and *k_g_* in the model, is shown for each of the 12 *P. polycephalum* organisms measured. Error bars for this ratio are standard deviations obtained from propagating errors of the measured parameters. This ratio is plotted against the average size of each organism, where the sizes are manually measured by summing up edge lengths *L* in images. Error bars for sizes are the standard deviations. The varied parameters produce different morphologies from more branched (e.g. organism 8) to less branched (e.g. organism 2). White traces show the time overlaid traces of the organism, meaning that multiple time point snapshots of the organism are pieced together to a single image that represents the locations the organism has traveled over. Black traces show the morphology of the organism at a single time point.

We modeled each organism using a traveling network model (22). The core traveling network model is a connected system that changes its location in space by rearranging its structure. The model assumes a binary tree with various connected nodes (Fig. 1d). The degree-one nodes (leaves), the nodes that are connected to only one other node, can either be in a “free” or “retracting” state. The free leaves of the network are allowed to undergo a number of actions such as growing, branching or switching. After switching, leaves undergo retraction until they encounter a degree-three node. These actions result in networks that travel away from their starting positions and explore space.

For each organism, we extracted quantities from time course imaging and converted them to corresponding model parameters (Fig. 1c) (see methods). To perform this conversion, we first determine the path, tail, and head of each organism’s trajectory through time. We define the path as the route connecting the network’s initial position to its final position, the tail as the oldest vertex in the network through time, and the head as the free leaf that moves along this path which never switches or retracts (Fig. 2A). Notably, the path, tail, and head are determined retrospectively, from timelapse imaging. Defining these concepts allows us to extract attributes for each organism that map to model parameters. Briefly, the branching rate parameter *k_b_* of the model is determined by identifying the number of branching events that occur along the path. Retraction speed was obtained by measuring the position of the tail throughout time and was used to determine the retraction rate parameter *k_r_* for the model. The growth rate parameter *k_g_* of the model is determined from a mathematical relation based on branching and movement. This relation is 2*k_b_* + *k_g_* = *f* ⋅ *k_r_*, where *f* ≤ 1 is the fraction of time the tail is retracting rather than paused (22). We observed that the position of the tail will have periods of time when it is moving versus stationary, which allows us to then compute this time fraction. The average size parameter in the model is determined by measuring the sizes (sum of edge lengths) of the organisms in multiple time lapse images. The branching angle parameter *α* is determined by measuring branching angles at the branching events along the path (Supplementary Figure 1).

**Figure 2:**
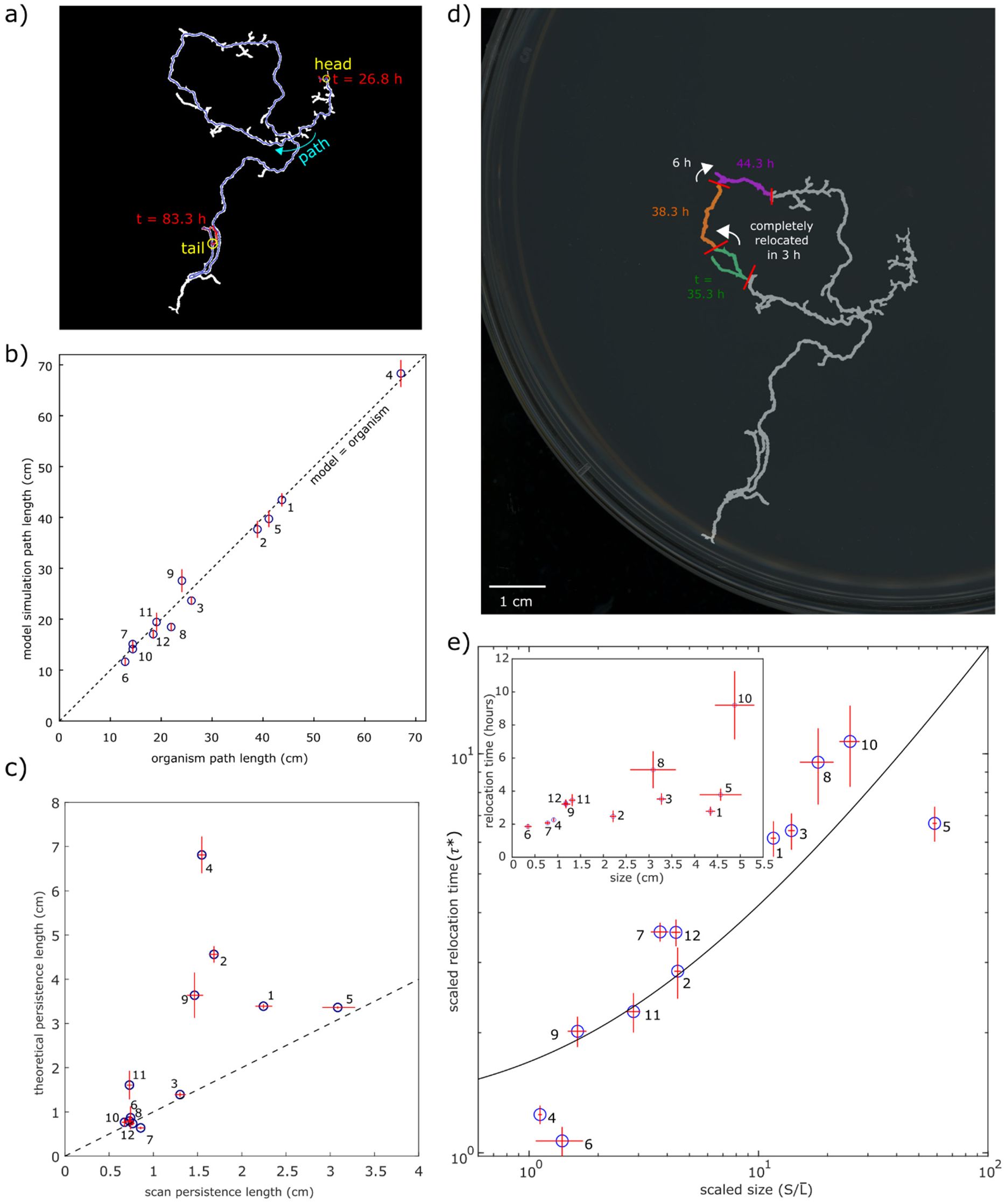
Results from a traveling network model map to emergent properties of *P. polycephalum*. a) The overlaid locations that one *P. polycephalum* has traveled through space, shown in white. The path is one single continuous trail from start to end within the white trace, formed by excluding the branches. The organism’s positions at the start and end of the time trace are shown in red, and the path is highlighted in blue. b) The path lengths of the organisms (n = 12) were compared to the path lengths of the corresponding simulations (99 simulations per organism). The diagonal line is the hypothetical perfect match between the organism and the model. Standard deviations are shown as error bars. c) Estimates of persistence length, derived from experimental coordinate data, against theoretical persistence length of the model. Error bars are standard errors, estimated from the confidence bounds of (experimental persistence length) and propagation of error from measured quantities (theoretical persistence length). Hypothetical line of agreement (dotted line) between the organism and the model is shown. d) *P. polycephalum* relocation time. Relocation is shown for an organism with overlaid images from multiple time points, where a few instances of organism locations after complete relocation are colored. For example, the organism was located at the position (green) at t = 35.3 h, and after 3.1 h, it had completely relocated and occupies a completely different position (dark orange). e) Relocation times were quantified for 12 *P. polycephalum* organisms. Inset shows relocation time against organism size. The main plot shows scaled relocation time (*τ*∗ = *N*_1_ ≈ 2*k_b_τ*) against scaled organism size (22). Solid line is the universal analytical relation (Eq. 4) derived from the model (not a fitting of the experimental data). Error bars are from propagating the error associated with each fundamental parameter (standard deviations) through the corresponding formulas.

These extracted quantities allow us to summarize the characteristics of each organism as a compact fundamental parameter set. Past work has shown that the organism can adopt different morphologies and behaviors in response to environmental conditions (16), or as they transition between different portions of the life cycle. In our experiments, stock organisms were maintained on oats, and a portion of the stock was harvested and inoculated onto nutrientless agar to initiate the experiment (see methods). This results in natural variability in both the size and state of the organisms upon initialization into otherwise identical experiments (Supplementary Videos 1 and 2). Across twelve organisms, we found that the *k_b_* parameter values ranged from 2.12 × 10^−3^ to 8.45 × 10^−3^, and the *k_g_* parameter values ranged from 0.54 to 0.91, where the rates are number of actions per 5 minutes (the duration between image acquisitions) for each leaf. The *k_b_* to *k_g_* ratio ranged from 2.33 × 10^−3^ to 1.14 × 10^−2^ (Fig. 1e, top). The average size ranged from 3.5 to 48.8 mm. In the model, a higher *k_b_* to *k_g_* results in a more branched morphology. Correspondingly, for two organisms of similar sizes we found that the organism with higher *k_b_* to *k_g_* ratio exhibits a more densely branched morphology, both at static snapshots and time-course history (Fig. 1e, bottom).

Rather than acting as a confounding factor, the substantial variability observed across these twelve organisms serves as a more rigorous test for the generalizability of our model. We next assessed whether a single traveling network model could capture the corresponding range of behaviors from this varied set of organisms.

### A traveling network model captures emergent properties of P. polycephalum

Next, we compared each *P. polycephalum* organism with its corresponding network model, where the input parameters are determined from each organism’s measurements. We assessed important *P. polycephalum* behaviors, including the route the organism takes as it travels through space and how structure and motion impact search at different resolutions. Notably, these behaviors are not directly specified in the model but emerge from the stochastic dynamics and fundamental parameters. To compare the organisms with the model, we relied on both analytical results that can be derived from the model and stochastic simulations of the model. For each organism we generated 99 corresponding simulation runs. We assessed the match between model results and experimental data for emergent properties that are important for foraging, including: path length, persistence length of the path, and relocation.

One important emergent property is the path length, which determines how far the organism can travel to forage for a food source. Here we measured path length for each organism and corresponding simulations (Supplementary Fig. 2). The path (Fig. 2a) is defined as the single continuous line that connects the organism’s starting and ending positions. Side branches may depart from the path, but for an acyclic traveling network, they will always retract back, leaving a single unique path. Figure 2b shows the relationship between measured and simulated path lengths. Organisms had path lengths varying by nearly an order of magnitude, and these were closely matched by simulations over the entire range, with 7/12 organisms falling within one standard deviation of the average from the simulated sets. The agreement between the path length of the organisms and corresponding models indicates that the traveling network model captures key information about how far these organisms travel through space.

The second emergent property we assessed is persistence length of the path. While the path length informs how far the organism travels, it does not indicate any changes in direction or shape of this path. Therefore, we extracted the persistence length, a metric indicating the characteristic length scale over which a random walk loses directional correlation, commonly applied to polymer chains (27). This provides us a metric of an average distance the organism travels before it changes direction. We apply this to the overall trajectory of the organism’s travels rather than a single time snapshot of the organism. We first marked coordinates along the path of each organism at a resolution of a few pixels, then calculated the persistence lengths by measuring the average change in the correlation of directional vectors spaced apart by different distances along the path (see methods). Persistence length from the model can be derived in terms of parameters as,

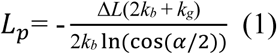

where Δ*L* is the change in edge length with each transition event (e.g. how much the length changes when one growth or one retraction event occurs) (22) and other parameters are as previously defined.

The organism and corresponding model persistence lengths are compared against each other in Figure 2c. There is general agreement between the organism and corresponding model persistence length, but some features stand out in this comparison. Persistence lengths for some organisms seem to systematically deviate above the model prediction (e.g. organisms 1, 2, 4, and 9). These organisms also tend to have lower *k_b_* to *k_g_* ratios (Fig. 1e), which points to a potential explanation for the difference. In the original traveling network model, growth is assumed to have perfect directional correlation (i.e. the only turning of the path happens through branching), whereas the organism’s path can change direction during growth alone, without branching (see, for example, the trajectory of organism 2 in figure 1e). This deviation is less pronounced for organisms that do relatively more branching. If capturing this aspect is desired, the model could be modified to lose directional correlation during elongation due to growth.

The third emergent property we assessed is a fundamental quantity of traveling networks, the relocation time, which is the amount of time that it takes for a network to occupy a completely new position (Fig. 2d). This quantity is associated with and unique to traveling networks; it does not exist for rooted networks which remain connected to their original location. We measured all the relocation events for each organism (Fig. 2d) and averaged these to obtain an average relocation timescale for each organism. Across organisms, the average relocation times varied from 0.37 hours (organism 6) to 1.84 hours (organism 10). The average relocation time tended to increase with increasing organism size (Fig. 2e, inset). Predictions from the model (22) suggest that the relocation time, *τ*, ultimately depends not directly on size, but on the number of leaves, *N*_1_, and the branching rate, *k_b_* according to,

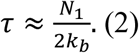

Previously, mathematical derivations demonstrated a relation between *N*_1_ and network size, *S*,

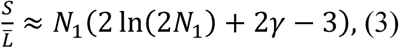

where *γ* is the Euler-Mascheroni constant, and *L̅* is the average edge length defined as 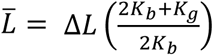

We can thus plot a “scaled relocation time”, *τ*∗, where *τ*∗ = *N*_1_ ≈ 2*k_b_τ*, against the scaled size, 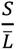, where the relation is as follows,

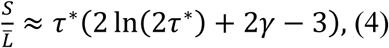

and evaluate whether data from the organisms matches predictions from the model.

Figure 2e shows the scaled relocation time vs scaled size for the twelve organisms and the model prediction, which can be plotted on these axes as a single analytical curve (Eq. 4). The model prediction captures the overall upward trend from the experimental data, which is especially meaningful since the model prediction is a universal equation, not an empirical fit to the data, and the trend is captured across organisms with over an order of magnitude differences in scaled relocation times and scaled sizes.

Taken together, we find that this simple traveling network model successfully captures behaviors related to the motion of *P. polycephalum* organisms by taking fundamental input parameters and predicting key emergent properties, including path length, persistence length, and relocation time.

### Edge width and avoidance help capture P. polycephalum search capabilities

Next, we quantify the amount of space the organism explores, which is important for the organism’s ability to forage. Evaluating the space covered by the organism is complicated by the irregular shape of the organism and its path through space and furthermore depends on the spatial resolution. To resolve these challenges, we applied a box-counting approach borrowed from the measurement of fractal dimensions (28). Box counting is conducted by placing the historical trajectory of the organism on different grids of varying box sizes, then counting the number of boxes occupied by the organism (Fig. 3a). Coverage of smaller boxes is an indication of finer search capabilities, while coverage of bigger boxes is an indication of coarser search capabilities. Therefore, coverage of a greater number of large boxes does not necessarily lead to coverage of a greater number of small boxes, and vice versa.

**Figure 3:**
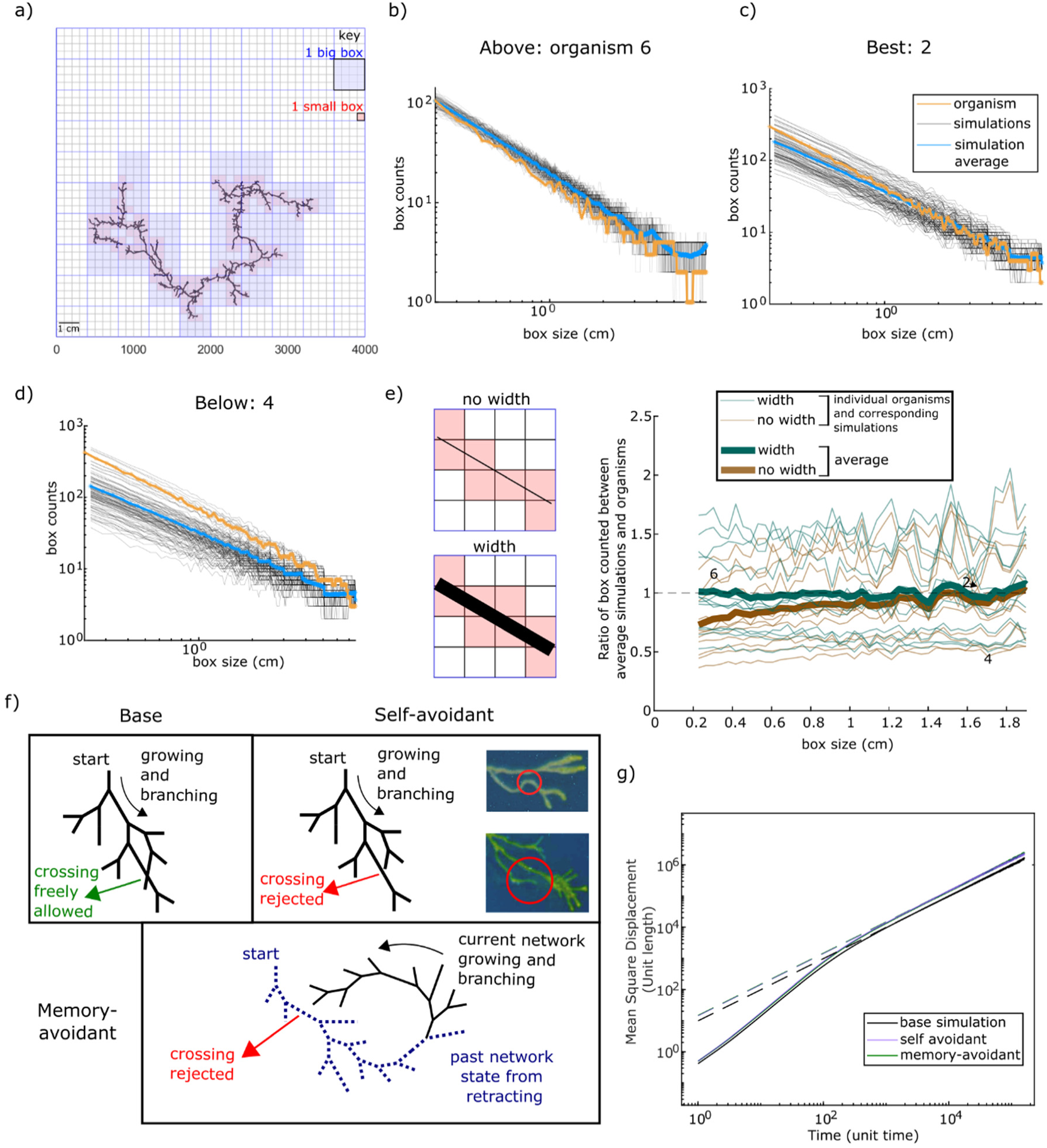
Search capabilities of *P. polycephalum* are captured by implementations of edge width and avoidance in the model. 3a) The amount of space covered by the *P. polycephalum* organism over time was quantified using a box-counting approach. The collection of bigger boxes (blue) illustrates the organism’s ability for coarser search and smaller boxes (red) illustrate finer search. 3b, c, d) *P. polycephalum*’s ability to collect boxes of different sizes is plotted along with the box sizes counted for the corresponding simulations. This is depicted for three organisms, 6, 2, and 4 for b, c, and d respectively. 3e) After adding width to the simulations, the boxes counted were quantified. The illustration (left) shows that the addition of width generally increases the number of boxes collected, particularly at small box sizes. The plot (right) shows the ratio in boxes counted between the simulations and corresponding organism, with a ratio of 1 representing the theoretical match of having equal amounts of boxes counted between organism and simulation. Numeric labels indicate the organism ID. 3f) Schematics of network avoidance behaviors. In the base form, the network is allowed to freely cross itself during growth and branching events. In self-avoidance, growth and branching at a leaf is rejected when encountering the organism itself. In memory-avoidance, the criteria are stricter, where growth and branching at a leaf is being rejected when encountering both the current position of the organism and areas the organism has already visited. 3g) Mean squared displacement over time for different avoidance models. Dashed lines are the tangential slopes at long time scales, representing the diffusive behavior of the simulated organism at later times.

We performed this box counting for the organisms and their corresponding 99 simulations, plotted for three representatives in figures 3b-e. First, when the box counts are plotted against the box size with log axes, we see that the typical slopes are between −1 and −2, indicating that the search history is more complex than a simple line but less than a complete space-filling dense pattern. We also see that the slope does not change considerably within a given organism, suggesting that the fractal characteristics of the organism do not vary substantially when viewing the organism at a small versus large scale. We see that the simulations of each organism show a range of box counts, due to the stochastic implementation. Specific organism box counts fall above (Fig. 3b), matching (Fig. 3c), or below (Fig. 3d) simulation averages, but data from the organisms is generally within the simulation envelope, with differences likely due to sampling the random process of motion.

In addition, we see a systematic trend that the number of boxes covered at small box sizes is generally less for the simulations than the corresponding organisms (Fig. 3b). This can be seen for specific examples, such as organisms 2 and 4 (Fig. 3c, d), and for the average ratio between the simulations and the corresponding organisms plotted across box sizes (Fig. 3e). This deviation indicates that the model looks too one-dimensional when zoomed in in comparison to the organism. This is due to the model consisting entirely of analytical lines without any width while the organism tubes have a small, but visible width. Therefore, a straightforward modification to the model is to simply give the edges of the simulated organism width, which should increase the number of small boxes counted (Fig. 3e, left). We implemented width into the simulation and found that this change eliminated the systematic underestimation at small box sizes without substantially changing the fit at larger box sizes, better capturing search behavior at the full range of length scales (Fig. 3e).

Another characteristic of *P. polycephalum* is avoidance (2,20). This avoidance was observed in *P. polycephalum* when it avoids revisiting areas it has previously explored through avoidance of its slime trails (2). In addition to avoiding past positions (Supplementary Video 3), the organism can also seemingly avoid the positions of its own current tubes (Fig. 3f, top right, circled in red) (Supplementary Video 4). We briefly show the possibility to incorporate this aspect into our model (Supplementary Video 5, see methods and code for implementation). To keep these two different forms of avoidance distinct, we refer to avoidance of previous positions as “memory-avoidance” and avoidance of currently occupied positions as “self-avoidance”. We show the distinction schematically compared to the original traveling network model (Fig. 3f).

We quantified the exploration capabilities of the three avoidant versions of this model: base (the original without avoidance introduced), self-avoidant, and memory-avoidant. Mean-squared displacement was used to quantify the space exploration of these networks (Fig. 3g). At shorter time scales, the network behavior is close to superdiffusive where the slope is much greater than one. In longer time scales, up to 1.5 × 10^5^ time units, the simulations for all three model versions have reached normal diffusive behavior with a slope closer to one. Both self-avoidant and memory-avoidant networks traversed more space than the base network as indicated by a slightly higher value for mean-squared displacement at large time scales. This implies that the avoidance capabilities of *P. polycephalum* may help it move faster through space.

We note that avoidance is not necessarily a strict rule for the organisms. From Reid *et al*. (2) and our own observations, the organism can still occasionally cross over locations it visited before. Furthermore, recent work has shown super-diffusive movement of *P. polycephalum* due to avoidance, which becomes more prominent as organism size increases (20). We also observe similar size-dependent avoidance that larger organisms are less likely to cross their previous positions than smaller organisms (Supplementary Figure 3). We note this only anecdotally due to the small number of observed crossing events, which limits the statistical significance of conclusions. The avoidant models could be easily modified to exhibit probabilistic avoidance, by including a tunable probability with which potential crossing events are accepted or rejected.

These results illustrate how the comparison between a traveling network model and experimental data can sharpen understanding by clarifying which elements are important for capturing system behaviors. For example, while consideration of network width was not important for a coarse-grain description of search, it was important for accurately capturing fine-scale search capabilities. Additionally, the model allows testing of features which may be difficult or impossible to control in experiments on organisms. For example, avoidance can take different forms that are readily implemented, tuned, and turned on and off in the model, revealing that avoidance enhances diffusivity.

## Discussion

In conclusion, here we presented the first explicit mapping of a traveling network framework onto a living organism, demonstrating that a simple traveling network model can accurately predict complex emergent behaviors for *P. polycephalum*, including the speed, directionality, search, and timescale of self-renewal.

A traveling network perspective provides a flexible framework to explore which elements are most important for capturing different system behaviors. Here we found that many aspects of *P. polycephalum* movement and search can be well captured with a simple traveling network model using just a few parameters. Here we showed how targeted addition of features such as avoidance and edge width can improve coarse- and fine-grained search. For *P. polycephalum*, other promising features to evaluate in future iterations of the model include higher-order branching, nonlinear edges, the formation and breaking of loops/cycles, and the splitting and merging of networks. Furthermore, this framework makes it possible to simulate sample sizes and time durations that remain logistically impractical in real-world experimentation.

While this organism is compelling in its own right, it represents a uniquely tractable example in the broader class of connected dynamic systems that travel through space. Such examples include the actin cytoskeleton (23,24), swarm robots (26), and human organizations (26). We do not expect each of these systems to conform to the exact model presented here; for example, actin networks can branch along edges rather than only at the leaves. Instead, we view this work as a guiding example of how to measure, map parameters, and model real traveling networks.

This work establishes how a traveling network model can be used to quantitatively predict key emergent properties from *P. polycephalum* behavior. We anticipate that a traveling network framework will be useful for studying the wide variety of systems that must dynamically alter their connected structures in order to move and explore.

## Materials and Methods

### Preparation of *Physarum polycephalum* plasmodia

*P. polycephalum* sclerotia (*Carolina Biological Supply*) was initially transferred onto a petri dish with 2% non-nutrient agar, along with oat flakes and water. *P. polycephalum* was allowed to grow from the sclerotia form to the plasmodial form in the dark. The plasmodial culture was maintained by periodically adding oat flakes and water, as well as transferring to a separate petri dish as necessary.

### Imaging of *P. polycephalum*

A small fragment of the plasmodia was then transferred to a separate larger petri dish (150 mm diameter) containing 2% non-nutrient agar for imaging under darkened conditions. A scanner (Epson Perfection V850 Pro) was used to take images every 5 minutes for 83.25 hours (999 images) or more, up to 143 hours. A black sheet was placed on top of the petri dish to create a black background. No additional food source (oat flakes) or water were added, to allow for free exploration. This was performed separately for a total of 12 *P. polycephalum* organisms.

### Measurements and analyses of *P. polycephalum* towards model input

After acquiring images for each *P. polycephalum* organism, measurements and analyses were performed on the images. Images were binarized in Ilastik (29), which were then overlaid to generate a full history of the organism.

The initial time points include a transient period where the organism begins moving and expanding from the initial placement. This transient period of the dynamics is assessed on a per-organism basis and was not included in the analysis.

Branching events were counted from this overlaid history, along with checking the original images for unclear occurrences. To account for differences between the model assumption (branching into a maximum of two) and the real organism (can branch into multiple), we counted branching into multiple as separate events (e.g. branching into two branches is considered one event, branching into three is considered two events, branching into four is considered three events, etc.).

Branching angles were then measured at each junction with a branching event on this overlaid history using ImageJ (30) to measure the angle between the tangential lines on the main branch following the path and the other most prominent branch.

Organism sizes were measured and estimated from selected original images in ImageJ, performed at a minimum interval of every 10 images to a maximum interval of every 100 images, dependent on organism. These were done by tracing freehand lines on the current positions of the organisms then summing the total length of those lines.

The retraction speeds were measured by viewing the original images and marking the coordinate of the tail end of the organism in ImageJ. This was performed at intervals of 10 images. Coordinate differences were computed at these intervals to obtain the retraction speeds. During this measurement, whether the tail is retracting or paused was also noted down. These overall measurements were performed on each organism.

### Conversion of measurements into model parameters

The aforementioned measurements were then converted into corresponding model parameters in the following ways. Each organism has its own unique set of parameters based on its corresponding measurements.

An average branching angle was obtained for each organism as the branching angle parameter for the model. The uncertainty in the branching angle was obtained from the standard deviation of the angular measurements (∼40-100 measured branching events per organism).

An average size was obtained for each organism corresponding to the equilibrium size parameter for the model. The uncertainty in size was obtained from the standard deviation of the size measurements (∼10-60 measurements per organism). The number of branching events was used to compute the branching parameter *k_b_* through dividing the number of branching events by the time duration of the images analyzed, then additionally dividing by two. The uncertainty in the branching parameter was obtained by dividing the square root of the counted number of branching events by time duration, then dividing this quotient by two (this follows from assuming branching events can be considered a Poisson process).

The retraction speed was determined by measuring the distances (in number of pixels) the organism retracts by for every 10 images, along with the times that the tail is retracting rather than paused. This distance is averaged over all the time points for which the tail end of the organism is retracting, converted to a value per image (5 minutes), and set to retraction rate *k_r_*. The uncertainty in the retraction rate is the standard deviation of these distance measurements. In the model simulation, rates were normalized by this retraction rate such that upper case *K_R_* is equal to one (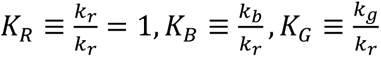, more details about the distinction between lower and upper case found in (22)). In the simulation, the unit length per action is set to one such that a length of one in the model simulation corresponds to the average retraction distance in pixels per image (or 5 minutes) of the particular organism. This means that a unit length of one in simulations of the model may correspond to a different real length from one organism to another, but this is simply a conversion difference on the relative scale of the simulated network versus the real organism. This conversion is accounted for in the comparisons when translating from the model to the organism and comparing between organisms. The growth parameter *K_G_* is then computed accordingly using 2*K_B_* + *K_G_* = *f* ⋅ *K_R_* (Cira, 2025). *f* is the fraction of time the tail is measured to be retracting rather than paused. Assuming *f* to follow a binomial distribution (tail moved vs did not move), the uncertainty in *f* was computed by inputting our measurements into the standard formula for standard deviation of a binomial distribution. The uncertainty in the growth parameter was obtained from propagating the error of the fundamental parameters through the above formula. This then gives us the main parameters *K_B_* and *K_G_* along with other parameters needed for the simulation.

### Measurement and Analyses of *P. polycephalum* towards emergent property comparisons

Several measurements were performed on the organism that were relevant for emergent property comparisons. These include path length, persistence length, and relocation time.

The path length was measured from the overlaid history by tracing a freehand line using ImageJ with a continuous line from the start to end point of the organism’s path.

The persistence length was measured from the overlaid history by drawing points along the path at a small distance resolution of every 4.18 pixels using Inkscape. The resulting drawn points were then measured in ImageJ to obtain a list of coordinates. Tangential vectors can be computed from these coordinate differences, where the tangential vector at a given point is the sum of the vector connecting to the previous point and the vector connecting to the next point. Angular differences can then be computed from these tangential vectors. Using cos 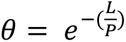 from polymer theory (31), a fit was performed between the measured data of *θ* denoting the angular difference and *L* the distance along the path between different angular pairs. An exponential curve was fitted using MATLAB, to arrive at a persistence length parameter *P* for each organism. Errors were from the 95% confidence bound fit output from the exponential fit, and standard errors were computed and plotted (Fig. 2c).

The relocation time was measured by viewing the original images. The initial position that the organism occupies was noted down. The next time point when the organism has completely occupied a new position was recorded. This was then repeated for subsequent time points. These time differences were averaged for each organism. Thus, an average relocation time was obtained for each organism then plotted (Fig. 2e). The uncertainty in the relocation time was obtained from the standard deviation of these measurements (6-62 relocations, depending on relocation time and duration of each dataset).

### Implementation of the traveling network model, model adjustments, and resulting analyses

The base version of the traveling network model (22) was simulated using MATLAB. Each organism has a unique set of model parameters from the measurements described above. 99 corresponding simulations were performed for each organism. Path length was computed from a custom MATLAB script that iteratively removes branches from the network structure such that only the path remains. The uncertainty of the path length was computed from the standard deviation of the path lengths of each set of 99 simulations. Persistence length was computed as previously derived from the model 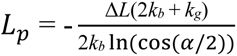 (22). Relocation time was computed from 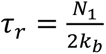 where *N*_1_ was calculated from *S* = *N*_1_(2 ln(2*N*_1_) + 2*γ* − 3) (22), where *S* is the average size of the network and *γ* is the Euler-Mascheroni constant. The scaled relocation time is *τ_r_* ∗ 2*k_b_* and the scaled size is 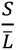 (22), where 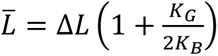. Both axes are then scaled by the retraction rate *k_r_*. Number of boxes covered by the simulation was computed through a custom MATLAB script that draws various-sized boxes then assesses based on coordinates whether any part of the network lies in each box. This was done for 99 simulations per organism then averaged.

Alternate versions of the traveling network model with width were implemented using a custom MATLAB script. A width value was added perpendicularly along the length of each edge of the network structure. The width value selected was based on a measurement of the visually largest tube width on each organism using ImageJ. Box counting for the width version of the model was performed as described above.

Alternate versions of the traveling network model with avoidance were implemented through modification of the base MATLAB script. Briefly, the simulation checks for line intersections with the current network position (tube-avoidance) or its past positions (memory-avoidance) whenever a growth or branching action occurs. In addition, a tunable parameter was also introduced that dictates the time criteria that decides whether the network is trapped or not. This allows the network to bypass a trapped state when no possible growth or branching actions are allowed without any line intersection (a temporary deviation from strict self-avoidance).

Memory-avoidance works similarly except that the past positions of the network are stored. A tunable parameter determines how many time steps into the past are stored for the previous network locations. Line intersection is checked against these past positions in addition to the current network position to determine whether growth or branching is allowed.

The base, self-avoidant, and memory-avoidant versions of the model were each simulated for 1.6e7 time steps. We used parameters of *K_G_* = 0.2, *K_B_* = 0.1, *S* = 100, *α* = 60°. In the data presented here, the network bypasses avoidance when it becomes trapped for 50 time steps. For the memory avoidance, the network avoids positions of the past 100 time steps. Mean squared displacements were computed by recording network centroid positions through time then computing the square of those position differences for corresponding time differences. The largest time step on the plot (Fig. 3g) is on the order of 10^5^ time steps even when the total run time is on the order of 10^7^ to allow for sufficient data points for mean square displacement to average over (i.e. there are many time intervals of 10^5^ within 1.6 × 10^7^ to give us many positions differences to average over). We note that long runtimes are required for simulating long time durations, and, while not used for the data presented here, a preliminary implementation indicates a version of the simulation written in Julia is ∼30x faster than the corresponding MATLAB version.

## Data and Code Availability

Raw video data for organisms 1-12 is available from Zenodo at 10.5281/zenodo.21908143. MATLAB scripts associated with this work are available at: https://github.com/CiraLab/AvoidantTravelingNetworks

## Acknowledgments

We thank members of the Cira Lab, and R. Schumann for stimulating discussions. N.J.C. acknowledges support from NIH grant R35GM157104. I.H.R.K. acknowledges support from NSF grants 2214020, and NIH grant 1R35GM163569.

