## Supplementary Information for "A traveling network model predicts emergent dynamics and search behavior from local remodeling in *Physarum polycephalum*"

### Supplementary Figures

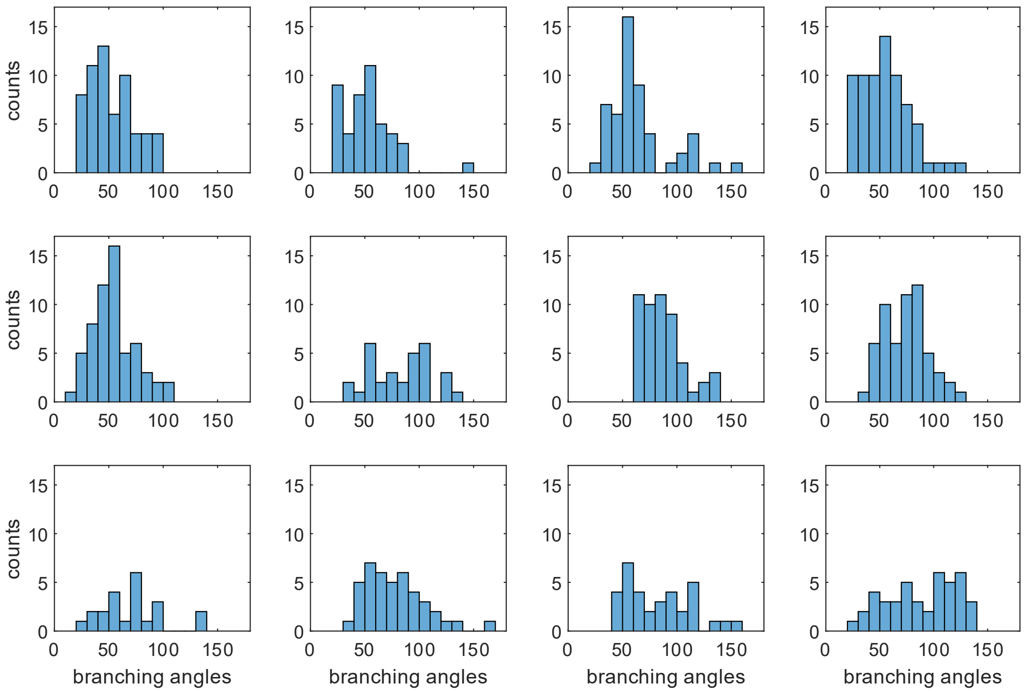


Supplementary Figure 1. Branching angle distribution for *P. polycephalum* organisms 1-12 (ordered left to right, then top to bottom), measured from time lapse traces of the imaged organisms.

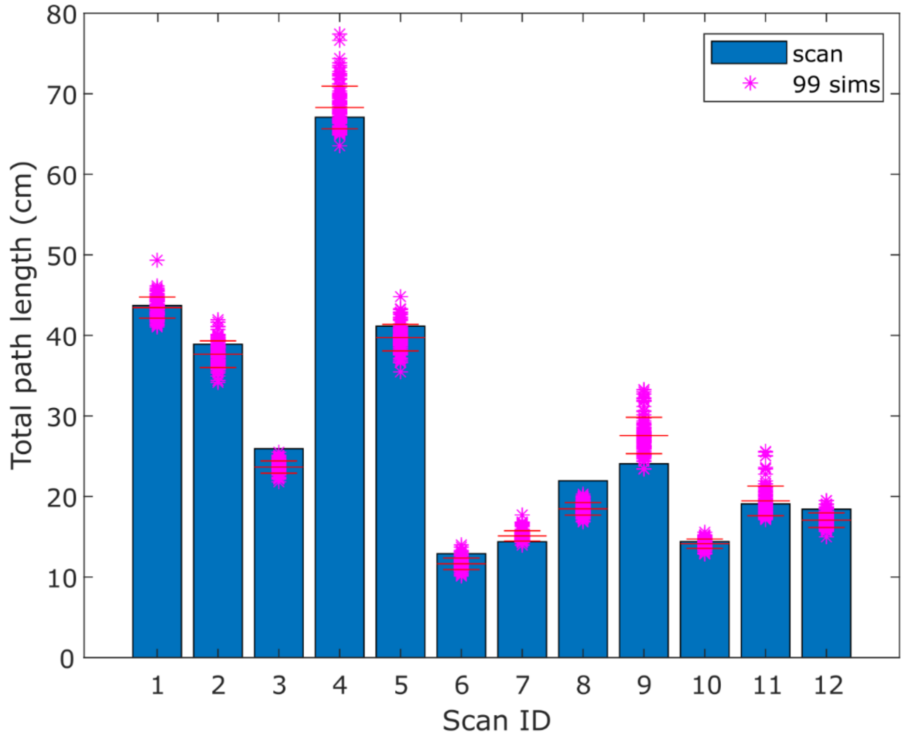


Supplementary Figure 2. Path length of each *P. polycephalum* organism compared with its corresponding 99 simulations. Each organism has its own separate corresponding 99 simulations, resulting in a total of 1188 simulations across 12 organisms. Each dot is the path length from an individual simulation. Horizontal lines indicate the standard deviation around the average.

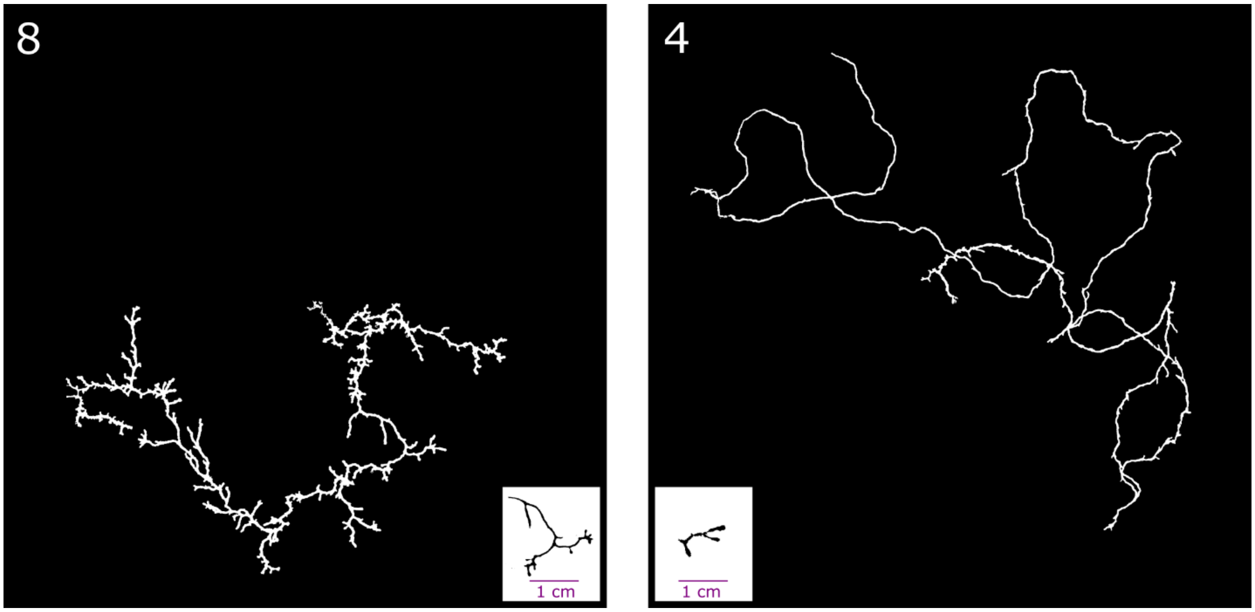


**Supplementary Figure 3.** Size-dependent avoidance for two different *P. polycephalum* organisms. The inset panels show a representative snapshot of each organism at a single time point to demonstrate the organism size (specific average size values plotted in Fig. 1e). The main panels show an overlaid trace of where the organism traveled through time. Here the larger organism, organism 8 (left, main) (average size of 3.09 cm) crosses itself fewer times as compared to the smaller organism, organism 4 (right, main) (average size of 0.91 cm). Previously, this size-dependent avoidance was observed when placing different sized organisms together and observing their avoidance interaction (20), and here we note how this applies to the trajectory of a single organism by itself over time. However, our observations of size-dependent avoidance are somewhat anecdotal, since most of the organisms do not cross their own trace more than once, if at all during the time scale of observation, restricting us from making robust comparison of size vs crossings for each organism.

**Supplementary Video 1.** Video of example *P. polycephalum* organism time trace (organism #4).

**Supplementary Video 2.** Video of example *P. polycephalum* organism time trace (organism #8).

**Supplementary Video 3.** Memory-avoidance of *P. polycephalum.* Example of *P. polycephalum* avoiding locations it has traveled to before.

**Supplementary Video 4.** Self-avoidance of *P. polycephalum.* Example of *P. polycephalum* avoiding the position of its own tubes.

**Supplementary Video 5.** Simulation of a self-avoidant traveling network simulation.
